# Long-read RNA sequencing identifies region- and sex-specific C57BL/6J mouse brain mRNA isoform expression and usage

**DOI:** 10.1101/2024.01.11.575219

**Authors:** Emma F. Jones, Timothy C. Howton, Victoria L. Flanary, Amanda D. Clark, Brittany N. Lasseigne

## Abstract

Alternative splicing (AS) contributes to the biological heterogeneity between species, sexes, tissues, and cell types. Many diseases are either caused by alterations in AS or by alterations to AS. Therefore, measuring AS accurately and efficiently is critical for assessing molecular phenotypes, including those associated with disease. Long-read sequencing enables more accurate quantification of differentially spliced isoform expression than short-read sequencing approaches, and third-generation platforms facilitate high-throughput experiments. To assess differences in AS across the cerebellum, cortex, hippocampus, and striatum by sex, we generated and analyzed Oxford Nanopore Technologies (ONT) long-read RNA sequencing (lrRNA-Seq) C57BL/6J mouse brain cDNA libraries. From >85 million reads that passed quality control metrics, we calculated differential gene expression (DGE), differential transcript expression (DTE), and differential transcript usage (DTU) across brain regions and by sex. We found significant DGE, DTE, and DTU across brain regions and that the cerebellum had the most differences compared to the other three regions. Additionally, we found region-specific differential splicing between sexes, with the most sex differences in DTU in the cortex and no DTU in the hippocampus. We also report on two distinct patterns of sex DTU we observed, sex-divergent and sex-specific, that could potentially help explain sex differences in the prevalence and prognosis of various neurological and psychiatric disorders in future studies. Finally, we built a Shiny web application for researchers to explore the data further. Our study provides a resource for the community; it underscores the importance of AS in biological heterogeneity and the utility of long-read sequencing to better understand AS in the brain.

## Introduction

Alternative splicing (AS) of preRNAs to mRNAs can result in multiple transcript isoforms and proteins from a single gene. This process contributes to the biological heterogeneity between species (1), sexes (2), tissues (3,4), and cell types (5). Notably, AS is more abundant in the brain, and the brain has more tissue-specific transcript isoforms than other tissues (3). AS is associated with many psychiatric and neurological disorders (e.g., Autism Spectrum Disorder (ASD), schizophrenia, and epilepsy (6,7)). Furthermore, many psychiatric and neurological disorders differ in prevalence by sex (8,9). For example, ASD, more common in males, has been linked to multiple genetic changes, including disordered splicing (10,11). However, as biomedical research has historically failed to study sex as a biological variable (12), there is still a need to quantify AS in the brain by sex accurately.

Recent advances in third-generation long-read sequencing technologies (i.e., Pacific Biosciences and Oxford Nanopore Technologies - ONT) enable high-throughput sequencing of complete mRNA transcripts to more rigorously determine the expressed transcript isoforms in a given sample compared to short-read (i.e., next- or second-generation) sequencing approaches. The resulting “long reads” can measure novel transcripts missed with prior studies and reveal extensive isoform-level diversity. For example, Clark et al. applied long-read sequencing to the human psychiatric risk gene *CACNA1C* and discovered 38 novel exons and 241 novel transcripts (13). While short-read gene expression AS data analysis can include calculating the percent spliced-in of exons or the splice junctions for a given gene, long reads enable researchers to quantify splicing across entire transcripts directly. Differential transcript usage (DTU), sometimes referred to as differential isoform usage, quantifies changes in transcript expression as a fraction of the overall expression of a particular gene, complementing differential gene expression (DGE) and differential transcript expression (DTE) analyses (14). Recently, researchers identified six candidate genes with novel DTU events in a schizophrenia cohort and developed a method to stratify patient populations using multi-gene DTU patterns (15), exemplifying that DTU can identify biologically relevant information in heterogeneous patient populations. These studies underscore how long-read sequencing approaches paired with novel analytical frameworks can identify and quantify AS patterns in the brain.

Due to known sex biases in healthy brain gene expression (2) and brain-related disease phenotypes (8,9), we studied AS across brain regions and sexes. Thus, we sequenced the cDNA from C57BL/6J mouse cerebellum, cortex, hippocampus, and striatum RNA for each sex (n = 5 each) using ONT and calculated DGE, DTE, and DTU between conditions. We generated over 85 million reads passing quality control metrics. We observed that the brain region with the highest DGE, DTE, and DTU is the cerebellum and that the most sex differences were in the cortex. We also built a web application hosting our data for use by the scientific community.

## Results

### Long-read RNA-Seq profiles across four mouse brain regions identified potentially novel genes and transcripts

We sequenced cDNA synthesized from total mRNA from the cerebellum, cortex, hippocampus, and striatum of 20-week-old male and female (n = 5 each) C57BL/6J mice using an ONT GridION device (**Figure 1A**). We obtained 85,909,493 reads passing quality control metrics (**Methods**), with each brain region receiving at least 16 million reads across the ten samples for that region (**Figure 1C**). The hippocampus had the lowest number of total reads (n = 16,739,487), potentially due to our reduced starting material as it is smaller than the other brain regions we assayed. We aligned and quantified our data using the nf-core (16) nanoseq pipeline and Bambu (17), a tool for performing machine-learning-based transcript discovery and quantification of long-read RNA-sequencing data with high precision and recall (18). When visualizing our samples based on variance-stabilization transformed (VST) gene counts by principal component analysis (PCA), samples are separated by tissue (**Figure 1B**). The difference in cerebellum samples to all other brain region samples drove the greatest gene expression variation in the data set (PC1, 33% of the total variance, **Figure 1B**).

**Figure 1.**
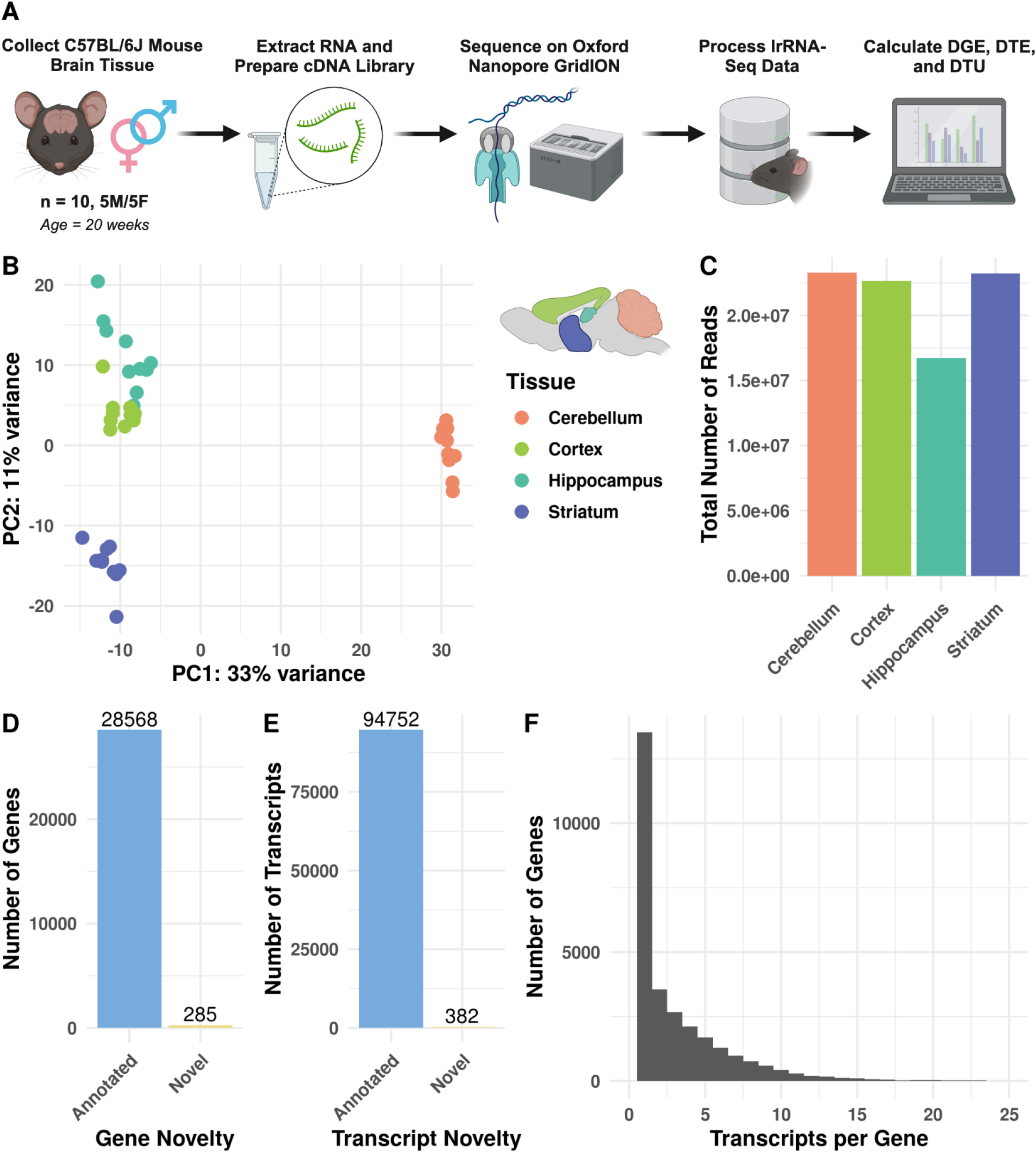
Long-read Nanopore RNA sequencing across four mouse brain regions. (**A**) Overview of the study design. (**B**) PCA plot (PCs 1 and 2) of VST gene counts. Here, we colored samples by brain region. (**C**) Bar graph of the total number of long reads sequenced for each tissue. (**D** and **E**) Bar graphs of the number of novel and annotated genes and transcripts. (**F**) Histogram of the transcript counts per gene, truncated past 25 transcripts per gene. **Supplementary File 2** includes the numbers of all transcripts measured for each gene.

We next determined any potentially novel genes and transcripts we captured with Bambu(17). We identified 285 genes and 382 transcripts not previously annotated in mouse GENCODE release M31 (**Figures 1D and 1E**) and considered them “novel.” These 382 potentially novel transcripts correspond to 354 unique genes. Of the 382 novel transcripts, 309 (81%) transcripts belonged to novel genes, and 73 (19%) belonged to previously annotated genes (**Supplementary File 1**). Interestingly, when we examined the expression distributions of novel compared to annotated transcripts, novel transcripts were expressed significantly more than annotated transcripts (Wilcoxon rank sum test, p = 1.570014e-113, 95% CI [2.80 4.15], **Supplementary Figure 1**), potentially due to the stringent expression cutoffs Bambu has to identify novel transcripts. Of these novel transcripts, 279 out of 382 (79%) had a mean counts per million (CPM) of at least one across all samples. However, all genes had a mean of 3.2 transcripts, while novel genes had a mean of 1.1 transcripts, though a subset of all genes (n = 76) had over 25 transcripts expressed (**Figure 1F, Supplementary File 2**). Two long non-coding RNA (lncRNA) genes, *Gas5* and *Pvt1*, had the most transcripts (149 and 129, respectively). In short, we generated a lrRNA-seq dataset for four brain regions and both sexes of C57BL/6J mice, in which we identified potentially novel genes, transcripts, and patterns of gene expression variance across mouse brain regions.

### Differential gene expression and differential transcript expression and usage identified across brain regions

We calculated DGE and DTE using the R package DESeq2 (19). We found 8,055 (Wald test with Benjamini-Hochberg (BH) correction p < 0.05) pairwise brain region DGE events involving 3,546 unique genes (**Figure 2A-D**), where the cerebellum, compared to the striatum, had the most DGE (n = 2,229, Wald test with BH correction p < 0.05), and the cortex, compared to the hippocampus, had the least DGE (n = 349, Wald test with BH correction p < 0.05) (**Figure 2B**). Consistent with our PCA (**Figure 1B**), each brain region compared to the cerebellum had the most DGE, with 920 genes consistently differentially expressed in the cerebellum compared to the other regions (**Figure 2B**). We calculated DTE for each expressed transcript, and we considered a gene to have DTE if it had at least one transcript with differential expression for that comparison (**Figure 2C-D**). We identified 11,138 DTE events (Wald test with BH correction p < 0.05) associated with 4,126 unique DTE genes (**Figure 2D**). Unlike DGE, the greatest difference in DTE genes was between the cerebellum and cortex (n = 2,620, Wald test with BH correction p < 0.05), and the least was between the cortex and the hippocampus (n = 345, Wald test with BH correction p < 0.05) (**Figure 2D**).

**Figure 2.**
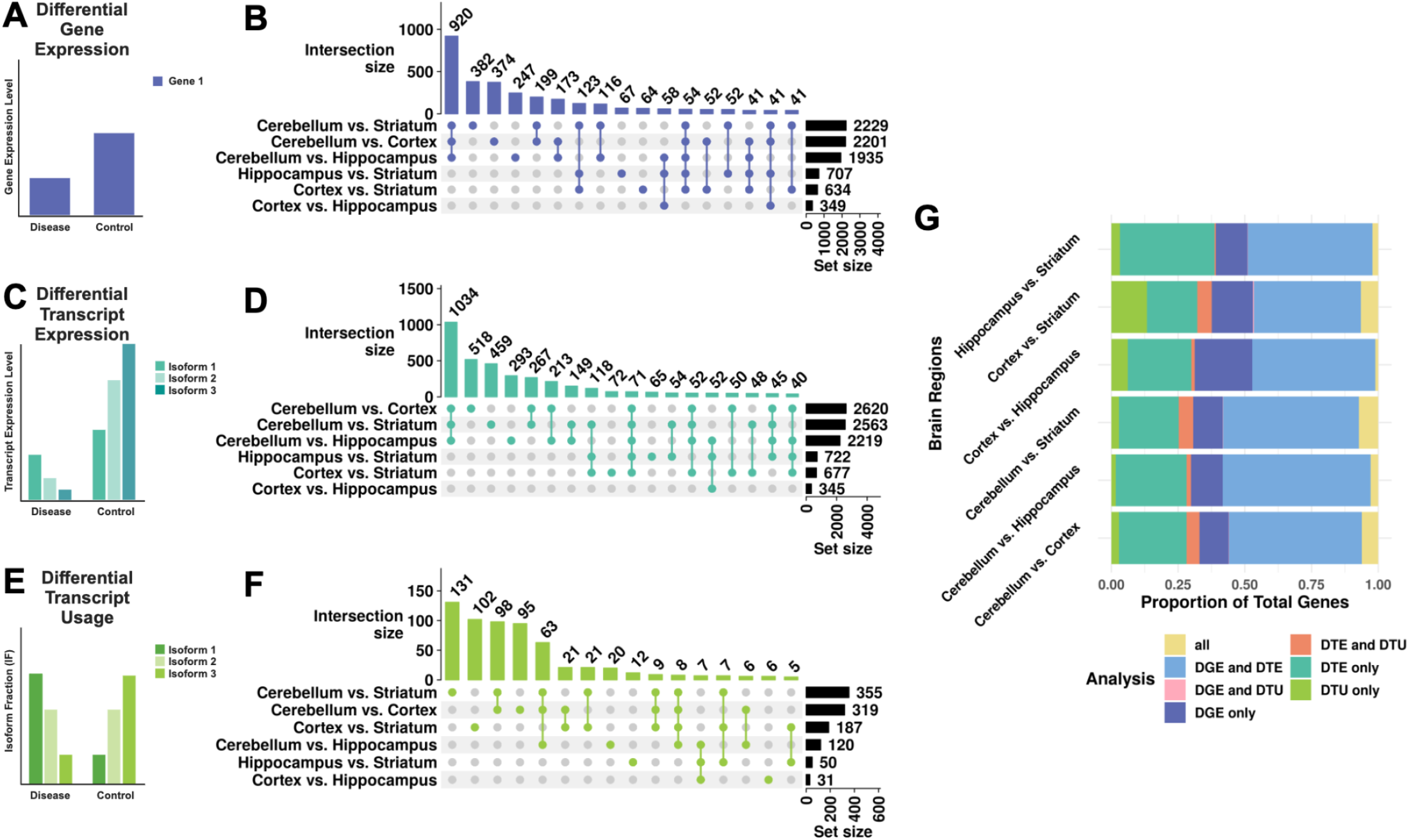
DGE, DTE, and DTU across pairwise brain region comparisons. Cartoon representation of a gene with three isoforms (actual genes may have more or fewer isoforms) exemplifying (**A**) differential gene expression (DGE - violet), (**C**) differential transcript expression (DTE - turquoise), (**E**) and differential transcript usage (DTU - green). UpSet plots of the overlap of genes with (**B**) DGE (Wald test with BH correction p < 0.05), (**D**) DTE (Wald test with BH correction p < 0.05), and (**F**) DTU (t-test with BH correction p < 0.05) between pairwise brain region comparisons. The bar plot above denotes intersection size, circles denote which comparisons have overlap, and the set size reflects the total number of genes with DTU for that comparison. For panels B and D, we omitted intersections of fewer than 40 genes from the chart for legibility. For panel F, we omitted intersections of fewer than five for legibility. (**G**) Stacked bar chart representing pairwise brain region comparison overlap across DGE, DTE, and DTU. Genes included in the chart must express at least two transcripts.

Next, we calculated DTU for each pair of brain regions using the DTU method SatuRn (20) with the R package IsoformSwitchAnalyzeR (21) (**Figure 2E, F**). Here, we considered a gene to be a DTU gene if it had a t-test statistic (calculated from the log-odds ratio and variance of the quasi-binomial generalized linear model) BH-corrected p-value < 0.05 for at least one of its transcripts where genes had at least two expressed transcripts (**Methods**). We analyzed DTU across brain regions and found 1,051 DTU events in 648 unique genes (**Figure 2F**). The most DTU genes were in the cerebellum compared to the striatum (n = 355, t-test with BH correction p < 0.05), and the least were in the cortex compared to the hippocampus (n = 31, t-test with BH correction p < 0.05) (**Figure 2F**). Consistent with our other analyses (65% for DGE and 71% for DTE), we identified the majority of DTU genes (66%) from comparisons including the cerebellum (**Figure 2B, D, F**). Interestingly, the number of DTU genes (n = 63, t-test with BH correction p < 0.05) shared across all three comparisons including the cerebellum was a smaller percentage (10%) of the total unique DTU genes (**Figure 2F**) than DGE (26%) or DTE (25%) (**Figure 2B, D**), suggesting that DTU analysis is less driven by the cerebellum. We also directly compared which genes were identified for each analysis (DGE, DTE, and DTU) that expressed at least 2 transcripts and qualified for DTU analysis. We found that DGE and DTE genes had the most overlap across comparisons, with a small proportion of significant genes for each comparison identified by all three methods (**Figure 2G**). We also performed functional enrichment analysis using gprofiler2 (22) of DGE, DTE, and DTU genes for all comparisons (**Supplementary Files 3-5**). For the cortex compared to the cerebellum DGE, DTE, and DTU genes, we found enrichment (Fisher’s exact test with g:SCS correction, p < 0.05) for 1742, 2431, and 54 terms. Strikingly, we found a much larger percentage of terms associated with the neuronal synapse in DTU (24/54, 44%; e.g., synapse, glutamatergic synapse, post-synapse, synaptic signaling, neuron-to-neuron synapse, and postsynaptic membrane) compared to DGE (50/1742, 2.9%) and DTE (77/2431, 3.2%). Because a larger proportion of DTU genes were enriched for pathways required for synaptic neurotransmission, this suggests that DTU potentially identifies biologically distinct molecular signatures from DGE and DTE. Overall, a pairwise comparison of DGE, DTE, and DTU between brain regions revealed marked heterogeneity for each analysis per comparison, with a greater overlap in DGE and DTE than in either analysis with DTU. This underscores that isoform usage may be masked when only considering differential expression, hiding biologically distinct molecular signatures.

### DTU sex differences are brain region-specific

Due to known sex biases in healthy brain gene expression (2) and in brain-related disease phenotypes (8,9), we asked if there were sex-biased DGE, DTE, or DTU events by brain region. First, we measured DTU across sexes, combining brain regions, and identified four genes with DTU: *Zfp862-ps, Gm10605, Shisa5*, and *Zfp324* (t-test with BH correction p < 0.05) (**Supplementary Figure 2**). *Zfp862-ps* and *Zfp324* are a pseudogene and gene, respectively, for zinc finger proteins that contain a DNA-binding domain. While pseudogenes have traditionally been considered non-coding, they have been shown to regulate other genes and form viable proteins (23,24). Notably, the human ortholog of *Shisa5, SHISA5*, has been previously identified as having sex-biased splicing in human brain white matter (2), in line with our finding of sex-biased splicing in mouse brain regions. Finally, *Gm10605* is a predicted lncRNA gene. We did not identify any of these genes in our within-brain region analyses, suggesting that for these genes, we were underpowered to identify DTU in each region alone.

We next calculated DGE, DTE, and DTU across sexes within each brain region (**Table 1, Figure 3**). We identified 23 region-specific genes with DTU by sex (analysis of deviance chi-squared test with BH correction p < 0.05): 14 in the cortex, seven in the striatum, and two in the cerebellum (**Table 1**). Despite documentation of phenotypic sex differences in the hippocampus (25), we did not find sex DTU in the hippocampus (**Figure 2C**). None of the 23 genes overlapped between brain regions, suggesting these sex differences may be brain region-specific. When we compared these DTU genes to DGE genes for each region, none overlapped (**Figure 3A-D**), and only three of 23 overlapped with DTE. Therefore, by analyzing DTU, we identified 20 additional genes with differential sex effects.

**Table 1.** Genes with brain-region-specific DTU across sexes.

| Brain Region | Genes with DTU across sexes |
| --- | --- |
| Cortex | 6430548M08Rik, **Anxa7**, *Plppr2*, *Sel1l*, *Zmiz2*, **Mtcl1**, *Kifap3*, *Leprot*, *Bcar1*, *Arhgap12*, *Washc3*, *Fbxw2*, *Bmal1*, *Lmtk3* |
| Striatum | *Fbxo25*, *Dhrs4*, **Rab28**, *Cacnb2*, *Cstpp1*, *Rsrc1*, *Celf2* |
| Cerebellum | *Camk2d*, *Srgn* |

Legend: Bolded genes are highlighted in the results and Figures 3 and 4.

**Figure 3.**
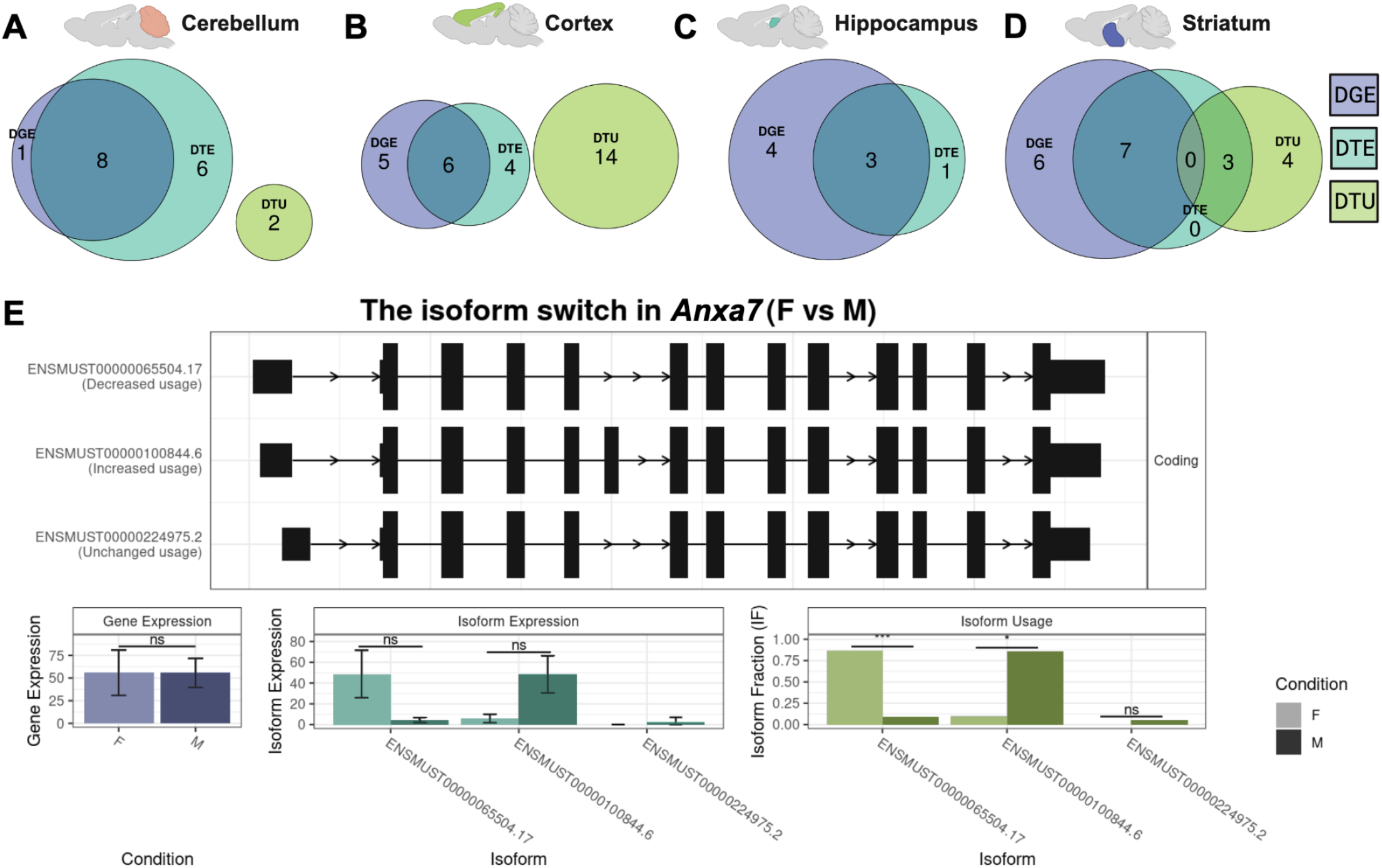
DGE, DTE, and DTU across sex within brain regions. (**A-D**) Euler diagrams represent the overlap of genes with significant DGE (Wald test with BH correction p < 0.05, purple), DTE (Wald test with BH correction p < 0.05, cyan), and DTU (analysis of deviance chi-squared test with BH correction p < 0.05, green). The brain regions represented are (**A**) cerebellum, (**B**) cortex, (**C**) hippocampus, and (**D**) striatum. (**E**) Switchplot displaying a transcript summary, gene expression, isoform expression, and isoform usage of the gene *Anxa7* across male (M; dark color) and female (F; light color) cerebral cortex. In the indicated comparison, ns denotes not significant, * denotes P < 0.05, ** denotes P < 0.01, and *** denotes P < 0.001.

We highlighted one of these sex-significant cortex DTU genes, *Anxa7*, for its many known connections to sex-associated phenotypes in humans (**Figure 3E**). Human *ANXA7* is a member of the annexin family, and humans express this gene in all tissues (26). *ANXA7* has multiple links to sex hormones; for example, *ANXA7* promoter activity is affected by estrogen and progesterone nuclear receptors (27). In addition, patients with schizophrenia express this gene lower than healthy controls (28). In our study, we measured three distinct *Anxa7* isoforms: ENMUST00000100844.6 (the Ensembl canonical transcript), ENMUST00000065504.7, and ENMUST00000224975.2 (**Figure 3E**). When we aggregate transcript expression, *Anxa7* does not have differential gene expression between males and females in any region. However, there was DTU of ENMUST00000065504.7 and ENMUST00000100844.6 across sex (**Figure 3E**). Males expressed ENMUST00000100844.6 (the only transcript that included exon 5) higher than females. Humans have a documented clinical variant of uncertain significance (gnomAD variant 10-75143086-T-A) in the conserved male-biased exon (29). Strikingly, 11/16 reported cases with this variant were in XY males and only 5/16 in XX females (29). In the alternatively spliced exon 5, multiple transcription factor binding sites exist, including for FOXO1, which is strongly sex-associated and a key transcription factor associated with early pregnancy (30). In summary, analysis of sex-significant DTU genes revealed differential isoform usage by sex within brain regions that would have otherwise been undetected by gene or transcript expression analyses, including genes with known sex-associated phenotypes.

### There are two main patterns of sexually dimorphic transcript usage: sex-divergent and sex-specific

In addition, we noticed distinct patterns in sex DTU genes expressing two transcripts (**Figure 4A-B**). First, we identified sex-divergent switches, i.e., sexually dimorphic transcript expression, where a single dominant transcript switch is in the opposite direction for both sexes (**Figure 4A**). We identified sex-divergent switches in *Mtcl1, Sel1l, 6430548M08Rik, Srgn*, and *Lmtk3*. For example, the sex-divergent gene *Mtcl1* has two transcripts, ENMUST00000086693.12 and ENMUST00000097291.10, where ENMUST00000086693.12 is dominant in males and ENMUST00000097291.10 in females (**Figure 4C**). *Mtcl1* codes for Microtubule Crosslinking Factor 1 and is expressed highly in the cerebellum in the literature and our dataset (26). Human *MTCL1* is known to be essential for the development of Purkinje neurons (31). Despite its connections to the cerebellum, we only saw DTU in *Mtcl1* by sex in the cortex. We also identified sex-specific isoform switches, i.e., where one sex expresses one isoform, but the other sex had almost equal expression of both isoforms (**Figure 4B**). We identified sex-specific isoform switches in *Rab28, Fbxo25, Leprot, Kifap3*, and *Plppr2. Rab28* (**Figure 4D**) has a female-specific isoform, ENMUST000000201422.4, which had approximately equal expression as the other isoform, ENMUST00000031011.12, in females, while ENMUST00000031011.12 was the only isoform expressed in males. *RAB28* is an essential gene for vision, and loss of function mutations in *RAB28* cause cone-rod dystrophy in humans (32,33). Thus, in addition to identifying significant differences in isoform usage between sexes, we also found distinct patterns of sex DTU gene expression, with sex-significant DTU genes showing either sex-divergent or sex-specific transcript expression.

**Figure 4.**
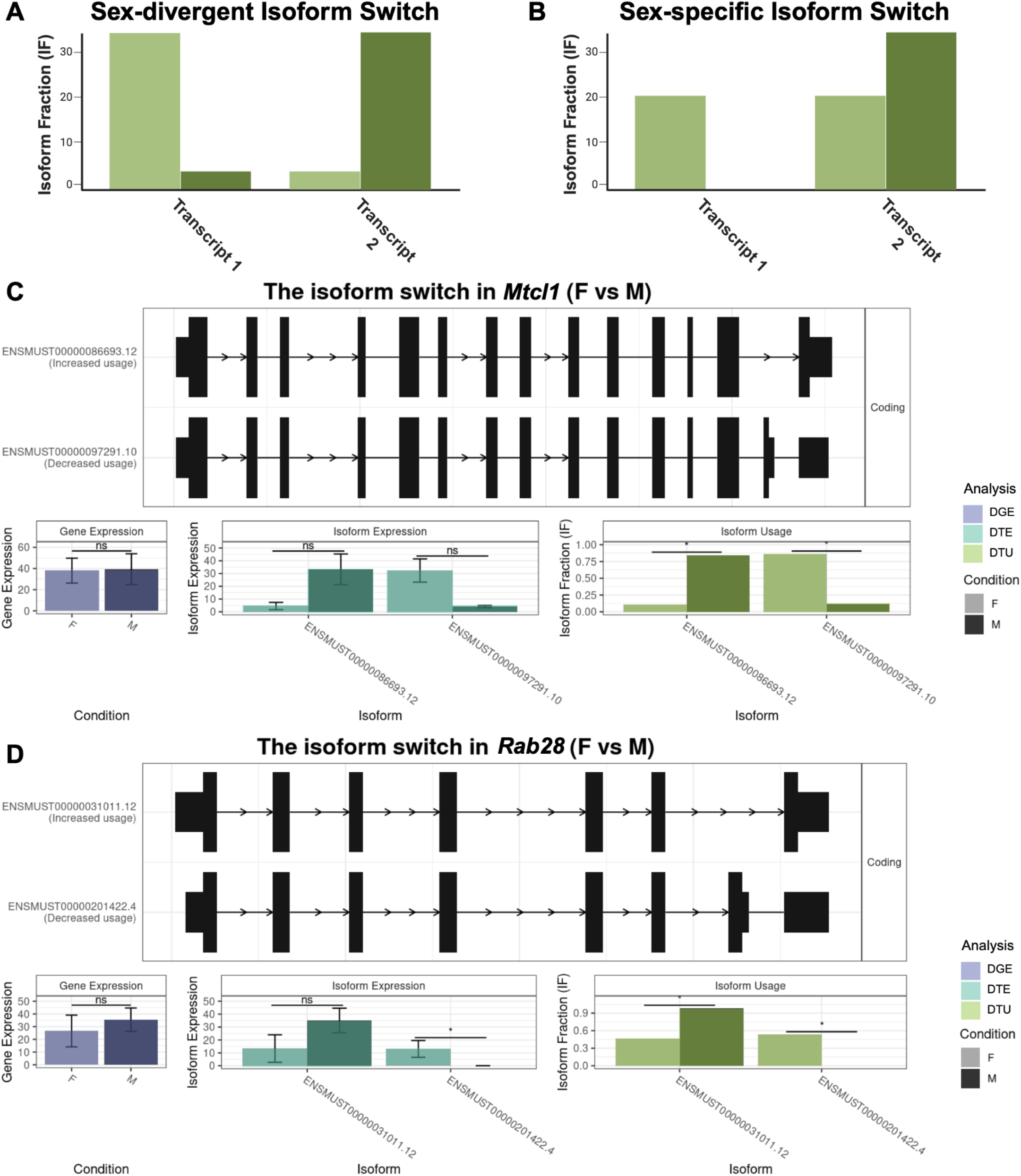
Sex-divergent and sex-specific DTU. (**A-B**) Representative cartoons exemplify two transcript expression patterns of isoform switching: sex-divergent (**A**) and sex-specific (**B**). (**C**) Switchplot displaying a transcript summary, DGE (Wald test with BH correction p < 0.05, purple), DTE (Wald test with BH correction p < 0.05, cyan), and DTU (analysis of deviance chi-squared test with BH correction p < 0.05, green) of the sex-divergent gene *Mtcl1* in the cortex between males (M; dark color) and females (F; light color). (**D**) Switchplot displaying a transcript summary, DGE (Wald test with BH correction p < 0.05, purple), DTE (Wald test with BH correction p < 0.05, cyan), and DTU (analysis of deviance chi-squared test with BH correction p < 0.05, green) of the sex-specific gene *Rab28* in the striatum between males (M; dark color) and females (F; light color). Please note that these plots do not display all possible transcript structures of this gene, only the ones measured in our dataset. In the indicated comparison, ns denotes not significant, and * denotes P < 0.05.

### A web application for visualizing DGE, DTE, and DTU in mouse brain lrRNA-seq data

Finally, we built an R Shiny application for our data set. Users may create custom gene expression heatmaps (**Figure 5A**) or examine switch plots for individual genes using the IsoformSwitchAnalyzeR package (**Figure 5B**). We also provide the option for users to download the intermediate gene expression and isoform switch test result data and plots directly. Our Shiny application has been made publicly available at https://lasseignelab.shinyapps.io/mouse_brain_iso_div/.

**Figure 5.**
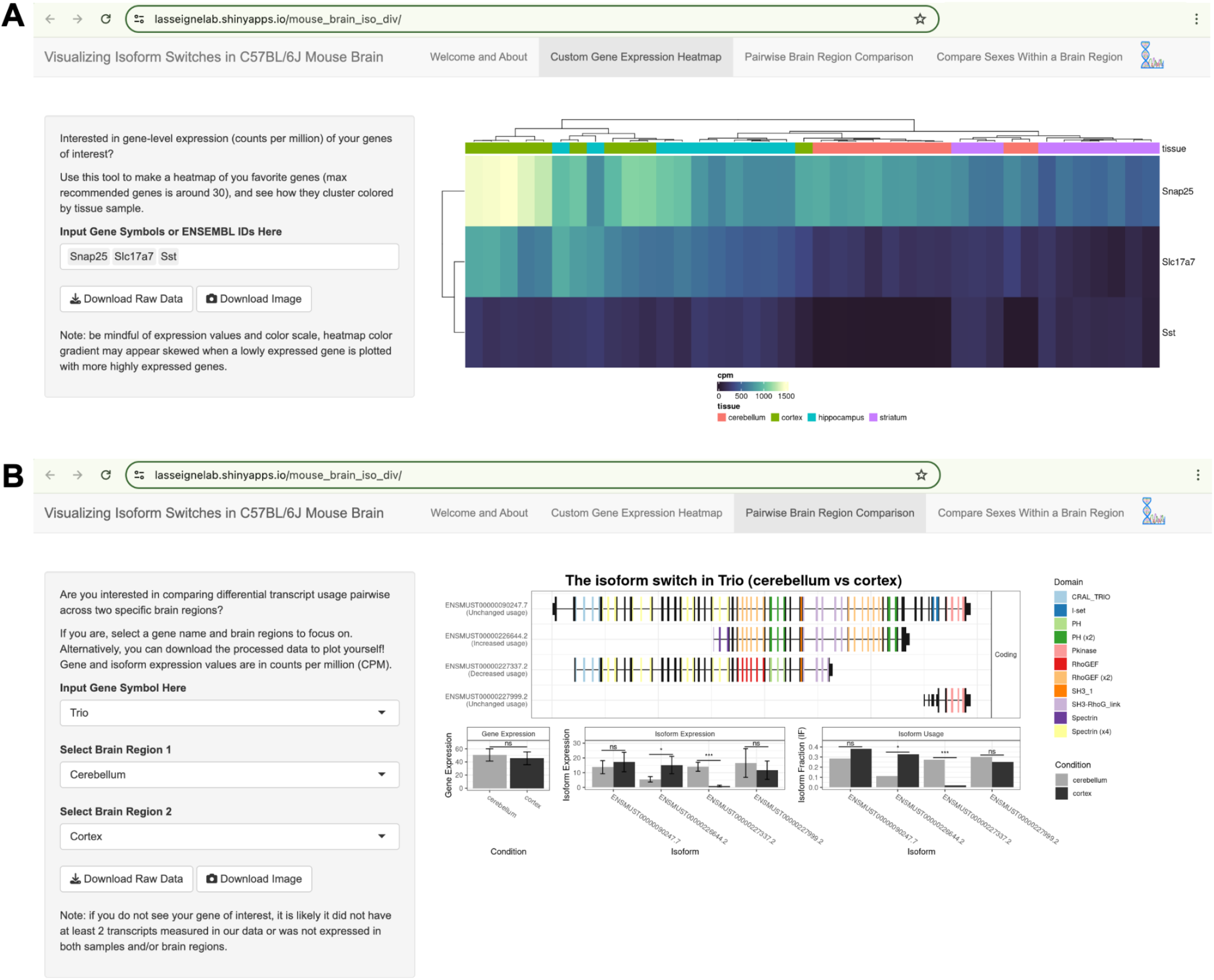
Shiny app presents a user-friendly interface for exploring our mouse brain dataset. Screenshots of our web application (**A**) The “Custom Gene Expression Heatmap” lets users examine and download the gene-level expression of any gene(s) of interest in our dataset. Users can also download the expression and isoform switch test result data to analyze further or download the plots as-is. (**B**) In the “Pairwise Brain Region Comparison” tab, users can visualize their gene of interest in pairwise brain region comparisons in real-time and download expression and isoform switch test result data and plots.

## Discussion

In summary, we produced a high-quality, publicly-available ONT lrRNA-Seq dataset across four brain regions from C57BL/6J mice, balanced for sex. We processed this data and identified 285 potentially novel genes and 382 novel transcripts, mostly (81%) associated with novel genes. We then calculated DGE, DTE, and DTU across brain regions and by sex. As expected, we identified DGE, DTE, and DTU between the four brain regions. The cerebellum had the most differences, potentially driven by cell type composition compared to the other three regions. Additionally, we found region-specific DTU between sexes, with the most differences in DTU in the cortex. We also report two distinct patterns of sex DTU in our data: sex-divergent and sex-specific. Finally, we built a Shiny web application for researchers to explore our lrRNA-Seq results.

Our study aligns with multiple prior studies identifying changes in isoform regulation across brain regions in mice (34–36) and humans (13,37–39). Additionally, we found the most differences in bulk DGE in the cerebellum, which agrees with other studies examining AS across multiple brain areas (40). For example, the gene with the highest DGE for all pairwise comparisons including the cerebellum is *Pcp2*, Purkinje cell protein 2. We suspect this reflects brain region-specific differences in cell type composition, as Purkinje neurons are unique to the cerebellum. However, confirmation of this hypothesis requires future studies at the single-cell level. Additionally, we found that some of these significant DTU genes across brain regions are known psychiatric risk genes (**Supplementary File 6**), potentially linking to region-specific differences in disease manifestation (41). We were not surprised by the low amount of DTU we observed across sexes when we grouped all brain regions because of the variability between different brain regions’ cell type compositions. Therefore, we also investigated AS across sexes within brain regions and found differences in the gene and transcript expression and usage of multiple brain-region-specific genes. Interestingly, we found the brain region with the most DTU by sex was the cortex, which is involved in high-level cognition. Many psychiatric phenotypes are associated with the cortex, and several of these are sex-biased in prevalence (e.g., ASD (8), schizophrenia (9), and major depressive disorder (42)). We also noticed that these DTU genes had two separate patterns of sex-significant transcript usage, either sex-divergent or sex-specific. These patterns demonstrate that while some transcripts are specific to one sex, others may shift in abundance between sexes, exemplifying nuanced sex differences.

We analyzed differences in gene expression on three fronts: DGE, DTE, and DTU, which together reveal more information on gene expression patterns by region and sex. While our work had many strengths, some limitations include using a bulk RNA-seq approach, read depth as a general limitation for transcript discovery, and sample number constraints, as well as using mice instead of human tissues for translation to human disease. Future work would benefit from single-cell resolution to determine the extent to which brain region differences stem from cell-type composition differences. Researchers could investigate this effect of cell-type composition through computational cell-type deconvolution, fluorescence-activated cell sorting (FACS), or new single-cell lrRNA-Seq methods, such as scISOr-Seq and scISO-Seq (43,44). While we sequenced an average of two million reads per sample and found 285 potentially novel genes and 382 novel transcripts, deeper sequencing depth may allow for greater novel isoform detection, as demonstrated by recently published AD data with extremely high-depth long-read sequencing (averaged 35.5 million reads per sample, discovered 3,394 new isoforms and 1,676 new gene bodies) (45). We reported on novel genes with any level of expression, and additional work is needed for our study and others to confirm these ORFs are actually novel genes and not sequencing bias or some other artifact. We attempted to reduce the number of false positives by using the stringent transcript quantification tool Bambu, which is specially designed for long-read sequencing data.

Furthermore, more samples may allow greater statistical power to detect smaller expression differences approaching significance with our current sample size. Intuitively, we would expect most DTU genes to have DTE, but not all DTE genes to have DTU. In our data, this assumption was not correct. Although SatuRn and DEXSeq use transcript expression information as the basis for their DTU analyses, these inconsistencies in significance between DESeq2 and SatuRn/DEXSeq may stem from using different models to calculate statistical significance (**Methods**). Therefore, it is possible that larger sample sizes and thus increased statistical power to detect significant differences in transcript expression and usage may result in the two methods agreeing more often for genes with DTU. Additionally, while mice and humans share many genetic similarities, our findings may not be directly translatable to humans. Surprisingly, we could not detect sex differences in alternatively spliced transcripts in the hippocampus, despite known sex differences in humans with hippocampal diseases (e.g., Alzheimer’s disease (46)). This may have been due to the sample input amount, sample numbers, species, or sequencing depth.

We aimed to examine and quantify differences across sexes and brain regions in C57BL/6J mouse brain tissue to better understand AS regulation. To our knowledge, this work is the first paper to use lrRNA-Seq to focus on brain-region-specific AS sex differences in the mammalian brain. We harnessed the power of lrRNA-Seq to investigate differences in AS with higher confidence than short-read and compared the results from three separate differential analyses. Here, we used novel sequencing technology to study sex as a biological variable, which is a necessary effort to resolve the long-standing practices of single-sex studies in preclinical biomedical research (12). In addition to publicly making all of our data and code available, we created an easily accessible web application for researchers to interact with the data. This research also serves as a launchpad for future directions involving additional time points, species, and disease contexts. Specifically, long-read spatial transcriptomics (47) and long-read ATAC (48) present opportunities for discerning patterns of AS and could be used to examine transcriptomic sex differences in isoform regulation at the spatial and epigenetic levels. Another future direction includes investigating classes of transcript diversity and structure (i.e., promoter usage and 3′ end choice) as done in ENCODE4 (49), but with an emphasis on studying differences across sexes in the brain. There is also a need to investigate sex differences in splicing across the lifespan, including early development (50,51) and aging (38). Finally, future research could combine long-read transcriptomics with measures of neuronal activity to discern the effects of AS on signal transmission across sexes (52). Our findings provide insight into sex differences in the mammalian brain, and the data produced by this research can serve as a useful resource for the scientific community.

## Materials and Methods

### Mouse sample collection and RNA isolation

We obtained flash-frozen hippocampus, striatum, cerebellum, and cortex C57BL/6J mouse brain tissues from The Jackson Laboratory (JAX #000664, age = 20 weeks) from five male and five female mice. The samples arrived on dry ice, and we stored them at -70°C upon arrival. For each sample, we transferred ∼30 mg of each brain region (or the entire brain region, in the case of hippocampus and striatum tissue) into an MP Biomedical Lysis D Matrix, 2ml tube (#6913500) containing 500μl of TRIzol reagent (Invitrogen #15596018) and lysed cells from each tissue on the FastPrep-24 5G bead beating grinder and lysis system (MP Biomedical #116005500). After lysis, we added 100μl of chloroform to the tube, centrifuged at 12,000×g for 15 minutes, and then transferred the clear top layer of the supernatant into a fresh tube. We next added an equivolume amount of isopropanol and centrifuged at 12,000×g for 10 minutes. We decanted the supernatant, washed the pellet twice with 75% ethanol, and resuspended the air-dried pellet in RNAse-free water. We incubated the final RNA product with TURBO DNase (Invitrogen #AM1907) for 30 minutes and assessed for RNA quality using a Qubit fluorometer and Agilent Fragment Analyzer. All RNA samples had an RNA quality number (RQN) score >7.

### Oxford Nanopore Technologies lrRNA-Seq library preparation

We processed RNA samples for nanopore sequencing using the PCR-cDNA Barcoding Kit (SQK-PCB111.24) according to manufacturer instructions and prepared libraries in equimolar amounts based on fragment length and concentration to make 15 fmol of cDNA library per flow cell. Because the barcoding kit only included 24 barcodes and we had 40 samples, we prepped and pooled two batches with 20 samples each. We loaded 11 μl of each pooled library with 1 μl Rapid Adapter T (12 μl total) onto 12 R9.4 flow cells (FLO-MIN106D). Because the Oxford Nanopore Technologies GRIDion (GRD-MK1) sequencing device can sequence five flow cells simultaneously, we sequenced these libraries in three separate sequencing runs for 72 hours each.

### Nanopore settings and software versions

We ran our nanopore with active channel selection turned on, a 1.5-hour pore scan frequency, a -170 mV initial bias voltage, and a -185 mV final bias voltage. We selected to have reserved pores off with high-accuracy base calling turned on. We used the following GridION software versions: MinKNOW 22.05.7, Bream 7.1.3, Configuration 5.1.5, Guppy 6.1.5, and MinKNOW Core 5.1.0.

### Raw sequencing data processing

We transferred demultiplexed FASTQ files to UAB’s supercomputer cluster, Cheaha, merged FASTQs passing a minimum Phred quality score of 9 for each sample and processed using the nf-core (16) nanoseq pipeline (https://doi.org/10.5281/zenodo.1400710) with the following options: version 2.0.1, protocol cDNA, flow cell FLO-MIN106, kit SQK-PCB109, skip_basecalling, skip_demultiplexing, skip_differential_analysis, profile cheaha, and a custom configuration file specifying nanoplot version 1.32.1. The packages we used for alignment and transcript quantification in this pipeline framework were Minimap2 version 2.17(53), samtools version 1.13 (54), and Bambu version 1.0.2 (17). We mapped reads using the GENCODE mm39 release M31 (available at: https://www.gencodegenes.org/mouse/) primary assembly genome and annotation. We retrieved transcript counts from the Bambu outputs of the nextflow results for further analysis.

### Data normalization

We processed and normalized data in R version 4.3.0 and RStudio version 2023.06.2+561. Because nanopore read lengths vary depending on the input cDNA length, we normalized by counts per million (CPM) instead of transcripts per million (TPM) since Bambu already accounts for length in its expression abundances. We calculated CPM by multiplying the number of read counts by 1 million and dividing by the sum of the total read counts for that sample. We found no outliers or batch effects by visual inspection when we performed principal component analysis (PCA).

### Differential gene and transcript expression analysis

For DGE and DTE analysis, we used the R package DESeq2 version 1.40.0 (19) using the negative binomial Wald test function. We considered a differentially expressed gene or transcript significance with a BH-adjusted p-value of less than 0.05 and an absolute log2 fold change >1.5 value. Therefore, we used three models:

- Region compared to another region (e.g., the cerebellum directly compared to the cortex)
- Sex within a region (e.g., female compared to male in the cerebellum)
- Sex across all regions (e.g., female compared to male)

We performed this analysis with gene-level counts for differential gene expression (DGE) and again with transcript-level counts for differential transcript expression (DTE). We then incorporated these results into the IsofrmSwitchAnalyseR switchList format for downstream plotting.

### Differential transcript usage analysis

We performed Differential Transcript Usage (DTU) analysis with the R package IsoformSwitchAnalyzeR package version 1.99.17 (21), using the satuRn version 1.8.0 (20) algorithm, and within brain regions, the DEXSeq version 1.46.0 (55) algorithm. Therefore, we used three models:

1. Region compared to another region (e.g., cerebellum compared to cortex) (satuRn)
2. Sex within a region (e.g., female compared to male in cerebellum) (DEXSeq)
3. Sex across all regions (e.g., female compared to male) (satuRn)

First, we created a switchAnalyzeRlist object with the importRdata function. We used the raw counts from Bambu (56) for the count matrix. For normalized isoform abundance values, we calculated CPM as described above. We used the IsoformSwitchAnalyzeR (21) package to remove genes that do not have more than one transcript and no gene expression minimum and proceeded with the satuRn (20) or DEXSeq (55) isoform switch tests. The satuRn isoform switch test uses a quasi-binomial generalized linear model to model transcript usage and calculates the posterior variance using an empirical Bayes procedure (20). Using this model, satuRn runs a t-test based on the model’s log-odds ratio estimates with the posterior variance and uses BH correction to reduce FDR (20). The DEXSeq isoform switch test uses a binomial generalized linear model and analyzes deviance for each “counting bin” based on a chi-squared likelihood ratio test (55). The IsoformSwitchAnalyzeR implementation of DEXSeq differs from other implementations of DEXSeq in that it uses full transcripts as the “counting bins” instead of exons so that it can detect DTU instead of only differential exon usage (21). Our significance filtering thresholds were an isoform switch q value < 0.05 and a differential isoform fraction (dIF) with an absolute value of at least 0.1, reflecting at least 10% change in isoform fraction across conditions. We calculated IF values as the isoform expression divided by total gene expression.

### Functional enrichment analysis

To infer pathways and diseases associated with the identified lists of significant genes with DGE/DTE/DTU, we performed a statistical enrichment analysis using gprofiler2 version 0.2.1 (22) with a custom set of background genes that passed filtering criteria (genes must have more than one transcript and be present in both conditions). We used the g:GOSt function, which uses a one-tailed Fisher’s exact test to obtain statistical probabilities for each term, and the g:SCS method for multiple testing correction. The default data sources for the gprofiler2 g:GOSt function include Gene Ontology (GO), Kyoto Encyclopedia of Genes and Genomes (KEGG), Reactome, Transfac, mirTarBase, CORUM, Human Protein Atlas (HPA), and Human Phenotype Ontology (HPO). We then saved the results in **Supplementary Files 3-5** and plotted these results, which passed our p-value threshold of < 0.05 for each comparison. When we compared the proportions of synaptic enrichment terms across analyses, we returned the number of terms that included the character string “synap”. We divided it by the total terms overall for that analysis.

### Comparison of DGE, DTE, and DTU

After determining which genes had DGE, DTE, and DTU for each condition tested, we created Euler diagrams and UpSet plots using the eulerr version 7.0.0 and ComplexHeatmap version 2.16.0 (57) packages, respectively, to visualize the overlap between these conditions. We identified genes with DTE by taking the unique list of gene IDs paired with transcripts identified as differentially expressed (adj p < 0.05) from DESeq2, where we only counted a gene with DTE in multiple transcripts once.

### Neurological disease phenotype gene sets

We compared three main gene lists to our significant DTU gene lists to known neurological disease risk genes. First, we compared against a recent set of Alzheimer’s Disease risk genes (58). Next, we compared against multi-disorder psychiatric risk genes from the Cross-Disorder Group of the Psychiatric Genomics Consortium (41). We listed psychiatric disorders if they have a posterior probability of association of above 0.9. Finally, we also compared active cases in UAB’s Center for Precision Animal Modeling (C-PAM).

To facilitate conversion between mouse and human genes, we converted the human neurological gene lists into mouse genes using the biomaRt Bioconductor package (59) in R. We then identified genes that were present in both DTU lists and neurological gene lists and reported them in **Supplementary File 6**.

### Protein domain family analysis

Following the package framework from the IsoformSwitchAnalyzeR package version 1.99.17, we extracted nucleotide and amino acid sequences from each gene’s open reading frame (ORF). Using those amino acid sequences as input, we ran the pfamscan.pl perl script with Perl 5 version 34 obtained from ftp://ftp.ebi.ac.uk/pub/databases/Pfam/ to identify known protein domains from the Protein family database (Pfam) (60). We incorporated these outputs into our R objects, and users can visualize select genes using our Shiny app.

## Supporting information

Supplementary Figures and File Descriptions

## List of Abbreviations

AS: Alternative splicing
ASD: Autism spectrum disorder
BH: Benjamini-Hochberg
CPM: Counts per million
dIF: Differential isoform fraction
DGE: Differential gene expression
DTE: Differential transcript expression
DTU: Differential transcript usage
lncRNA: Long non-coding RNA
lrRNA-Seq: Long-read RNA sequencing
ONT: Oxford Nanopore Technologies
ORF: Open reading frame
PCA: Principal component analysis
RQN: RNA quality number
TPM: Transcripts per million

## Availability of Data and Materials

The raw dataset supporting the conclusions of this article is available in the Gene Expression Omnibus (GEO) repository, with accession number GSE246705, https://www.ncbi.nlm.nih.gov/geo/query/acc.cgi?&acc=GSE246705

The docker images, intermediate datasets, and code to reproduce all analyses and results in this article are available in the following Zenodo repositories: Docker images - https://zenodo.org/records/10480924, intermediate data -https://zenodo.org/records/10381745, GitHub code - https://zenodo.org/records/10481313.

The code supporting the conclusions and for reproducing analyses of this article is available in the GitHub repository, https://github.com/lasseignelab/230227_EJ_MouseBrainIsoDiv.

The interactive web browser application associated with this manuscript is available at https://lasseignelab.shinyapps.io/mouse_brain_iso_div/.

## Authors’ Contributions

EFJ and BNL conceptualized the project. EFJ, TCH, and VLF collected and generated the sequencing data set. All analyses were coded and performed by EFJ. EFJ developed and deployed the web application. TCH, VLF, and ADC reviewed and validated the code. BNL and TCH provided supervision and project administration. BNL acquired funding. EFJ wrote the first draft. EFJ, TCH, VLF, ADC, and BNL reviewed and edited the manuscript. All authors read and approved the final manuscript.

## Acknowledgments

We acknowledge all current and past members of the Lasseigne Lab for their thoughtful feedback, especially Tabea M. Soelter, Jordan H. Whitlock, Vishal H. Oza, and Elizabeth J. Wilk. We would like to thank the UAB Biological Data Sciences (UAB-BDS) core for discussions during office hours and institutional support of the Cheaha configuration for nf-core pipelines and docker/singularity container documentation. All figures and cartoons were assembled with BioRender.

