## Supplementary Figures and File Descriptions for "Long-read RNA sequencing identifies region- and sex-specific C57BL/6J mouse brain mRNA isoform expression and usage"

**Supplementary Figure 1. Novel transcripts are expressed more highly than annotated transcripts.** Density plot showing expression levels (measured in counts per million, CPM) between the novel (blue) and annotated (yellow) transcripts. Density is measured by kernel density estimation. The two distributions were significantly different when we ran a Wilcoxon rank sum test, p = 1.570014e-113.

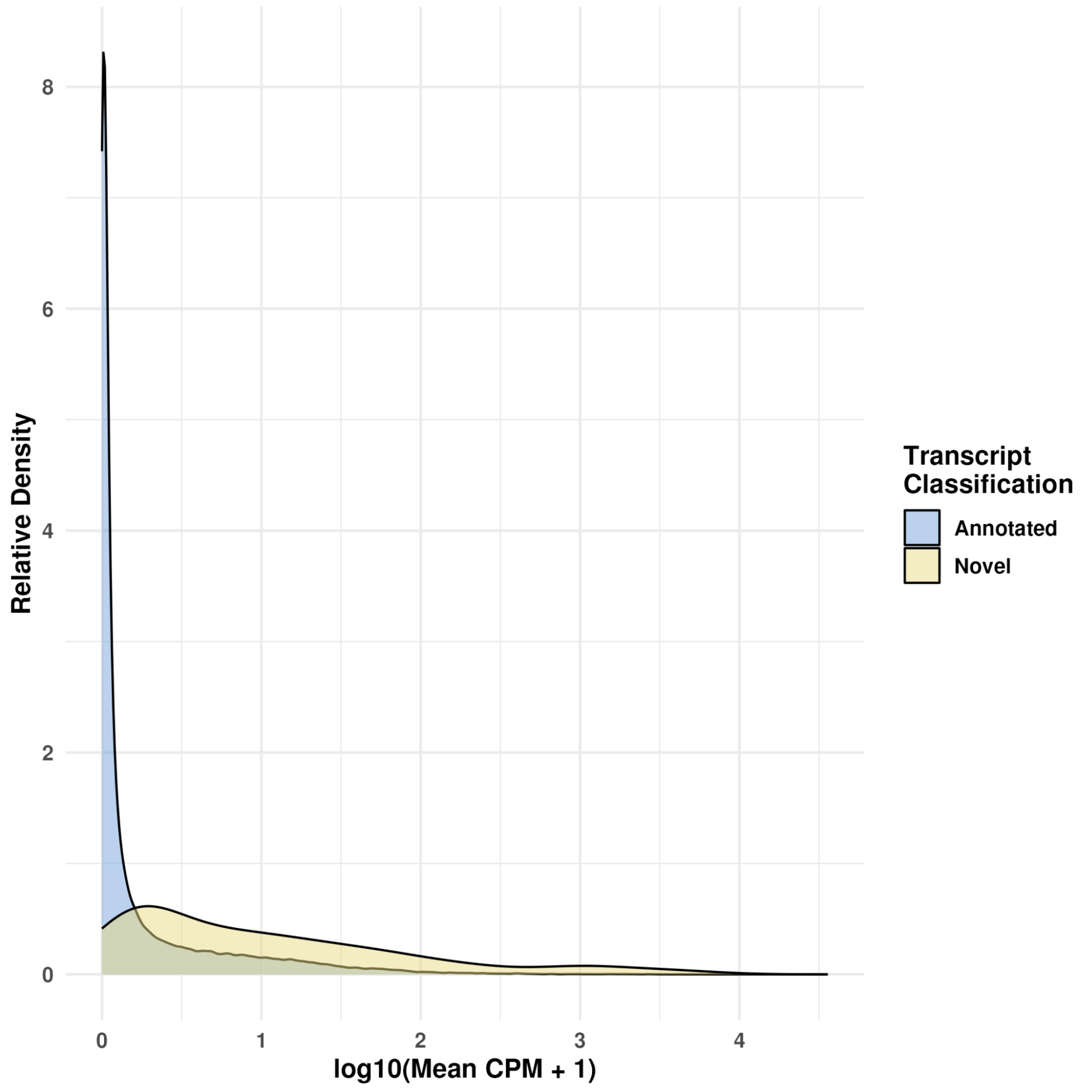

**Supplementary Figure 2. Genes with DTU across sex for the whole brain.** SwitchPlots showing transcript summary, gene expression, isoform expression, and isoform usage of the genes for all brain regions across males (M; dark gray) and females (F; light gray). Genes shown are *Zfp324* (**A**), *Zfp862-ps* (**B**), *Shisa5* (**C**), and unannotated gene or *Gm10605* (**D**).

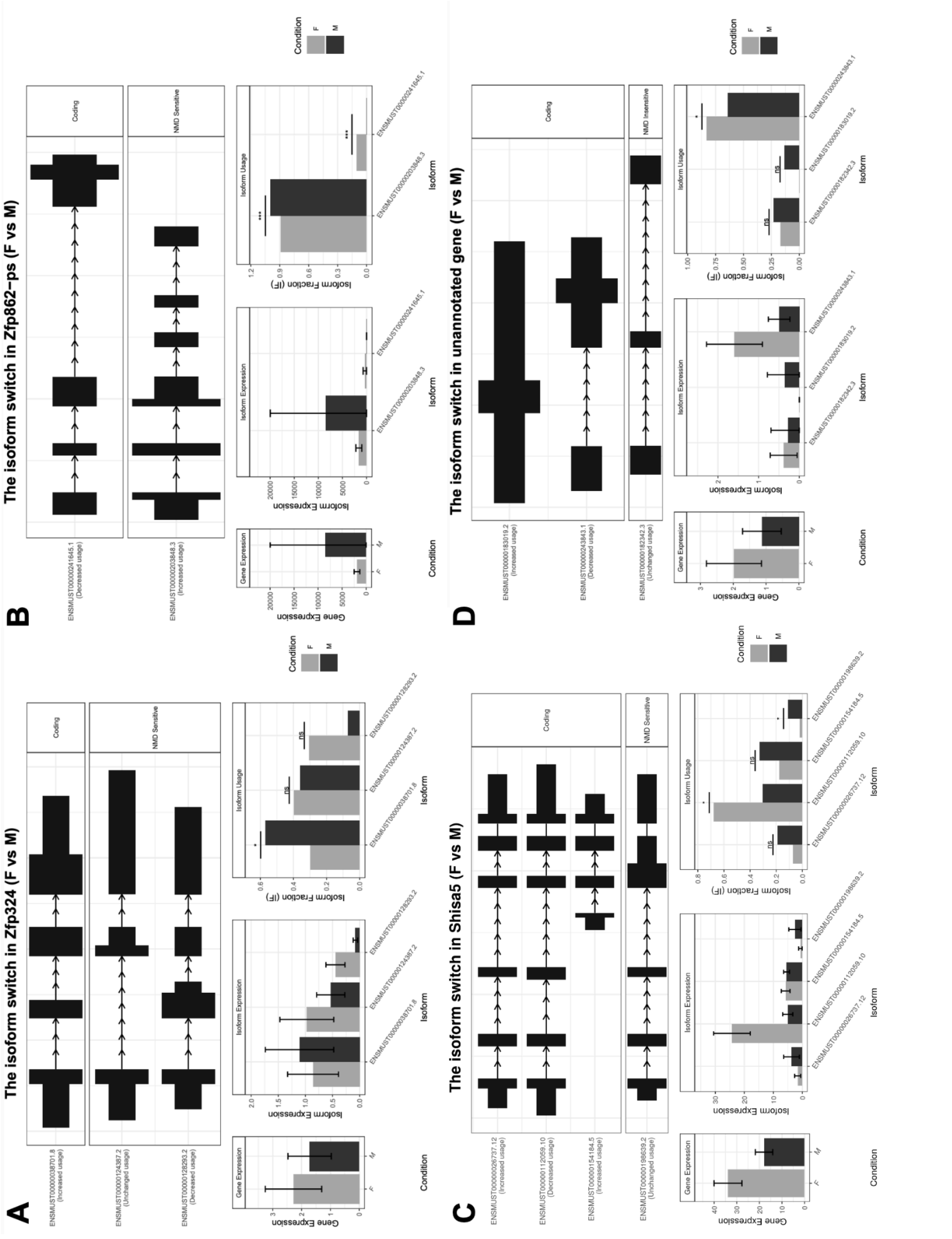

**Supplementary File 1. Novel transcripts information**
Legend: This supplementary file is a CSV with general information about the novel transcripts.
Columns as as follows:

- Location - which chromosome we found this gene on
- Start - the genomic start position of the novel gene
- Stop - the genomic end position of the novel gene
- Strand - the direction of transcription for the novel gene
- Gene_id - the gene identification number automatically assigned by Bambu
- Transcript_id - the transcript identification number automatically assigned by Bambu

Filename/location: 230227_EJ_MouseBrainIsoDiv/results/novel_txs/novel_txs_annotation.csv

**Supplementary File 2. Transcripts per gene table**
Legend: This supplementary file is a CSV with the number of all transcripts measured for each gene.
Columns are as follows:

- GENEID - Either ENSEMBL gene identification number, if available, or Bambu assigned identification number
- Transcript count - number of transcripts counted per gene with more than 0 counts

Filename/location: 230227_EJ_MouseBrainIsoDiv/results/all_txs/txs_per_gene.csv

**Supplementary File 3. DGE Functional enrichment results**
Legend: This supplementary file is a Microsoft Excel file with gprofiler results of DGE genes with a sheet for each comparison
Columns are as follows:

- Query - all results were processed as individual queries
- Significant - all terms in this table were kept if they had a significance of below 0.05
- P_value - p-value from Fisher's one-tailed test
- Term_size - number of genes in this term size
- Query_size - number of genes for this specific query
- Intersection_size - number of genes in the intersection between term and query
- Precision - statistical precision for this term
- Recall - statistical recall for this term
- Term_id - Identification number for this term
- Source - Data source for this term
- Term_name - Name for this term
- Effective_domain_size - Domain size for this term
- Source_order - Order for this term
- Parents - Any parent terms for this term

Filename/location: 230227_EJ_MouseBrainIsoDiv/results/gprofiler2/dge/

**Supplementary File 4. DTE Functional enrichment results**

Legend: This supplementary file is a Microsoft Excel file with gprofiler results of DTE genes with a sheet for each comparison
Columns are as follows:

- Query - all results were processed as individual queries
- Significant - all terms in this table were kept if they had a significance of below 0.05
- P_value - p-value from Fisher's one-tailed test
- Term_size - number of genes in this term size
- Query_size - number of genes for this specific query
- Intersection_size - number of genes in the intersection between term and query
- Precision - statistical precision for this term
- Recall - statistical recall for this term
- Term_id - Identification number for this term
- Source - Data source for this term
- Term_name - Name for this term
- Effective_domain_size - Domain size for this term
- Source_order - Order for this term
- Parents - Any parent terms for this term

Filename/location: 230227_EJ_MouseBrainIsoDiv/results/gprofiler2/dte/

**Supplementary File 5. DTU Functional enrichment results**
Legend: This supplementary file is a Microsoft Excel file with gprofiler results of DTU genes with a sheet for each comparison
Columns are as follows:

- Query - all results were processed as individual queries
- Significant - all terms in this table were kept if they had a significance of below 0.05
- P_value - p-value from Fisher's one-tailed test
- Term_size - number of genes in this term size
- Query_size - number of genes for this specific query
- Intersection_size - number of genes in the intersection between term and query
- Precision - statistical precision for this term
- Recall - statistical recall for this term
- Term_id - Identification number for this term
- Source - Data source for this term
- Term_name - Name for this term
- Effective_domain_size - Domain size for this term
- Source_order - Order for this term
- Parents - Any parent terms for this term

Filename/location: 230227_EJ_MouseBrainIsoDiv/results/gprofiler2/dtu/

**Supplementary File 6. DTU genes that are known neurological disease risk genes**
Legend: This supplementary file is an Excel file of neurological disease risk genes and which DTU design we used to identify them. We compared significant DTU genes to lists of Alzheimer's disease, cross-psychiatric disorder, and UAB's Center for Precision Animal Modeling case genes (C-PAM) to identify disease genes.

Filename/location: /230227_EJ_MouseBrainIsoDiv/results/psych_dtu_genes/
